# Cellular profiling identifies an early profibrotic alveolar type 2 cell signature in lung fibrosis

**DOI:** 10.1101/2025.05.22.655638

**Authors:** Ram P. Naikawadi, Alexey Bazarov, Michael Wax, Kaveh Boostanpour, Jasleen Kukreja, Michela Traglia, Ayushi Agrawal, Reuben Thomas, Mallar Bhattacharya, Paul J. Wolters

**Affiliations:** Department of Medicine, Division of Pulmonary, Critical Care, Allergy and Sleep Medicine, University of California, San Francisco, CA, USA; Department of Surgery, University of California, San Francisco, CA, USA; Gladstone Institute Bioinformatics Core, San Francisco, CA, USA

**Author notes:** Correspondence: Paul Wolters, University of California, San Francisco, 505 Parnassus Ave, Room M1090, San Francisco, CA 94143.

**Keywords:** Telomere, fibrosis, senescence, interstitial lung disease, Alveolar type 2 cells, p53

## Abstract

**Rationale:** Idiopathic pulmonary fibrosis (IPF) is a progressive, age-associated, lung disease characterized by short telomeres in alveolar type 2 (AT2) cells, epithelial remodeling, and fibrosis.

**Objectives:** This study investigated how telomere dysfunction in AT2 cells lacking Telomere Repeat Binding Factor 1 (TRF1) drives lung remodeling in SPC-creTRF1^flox/flox^ mice and its relevance to IPF.

**Methods:** Mouse model of telomere dysfunction was used to conditionally delete TRF1 in AT2 cells. SPC-creTRF1^flox/flox^ mouse lung epithelial cells were used to perform single cell RNA sequencing. AT2 cells from IPF lungs were analyzed by single cell RNA sequencing in an organoid model.

**Measurements and Main Results:** Single cell RNA-sequencing revealed distinct pathological AT2 cells enriched in DNA damage, senescence, oxidative stress, and pro-fibrotic genes, along with fewer “normal” AT2 cells and increased club cells in SPC-creTRF1^flox/flox^ mice. Pathological AT2 cells showed different early and late-stage gene signatures, with a prominent p53 signature at both time-points. Genetic deletion of p53 in SPC-creTRF1^flox/flox^ AT2 cells improved survival and prevented lung fibrosis. p53 deletion or inhibition improved organoid formation, surfactant protein C expression, and reduced pro-fibrotic gene expression in AT2 cells isolated from SPC-creTRF1^flox/flox^ mice or IPF lungs.

**Conclusions:** These data suggest that the DNA damage response to AT2 cell telomere dysfunction, driven by enhanced p53 activity, mediates early AT2 cell transdifferentiation and senescence, leading to epithelial cell remodeling and fibrosis and that reversing this reprogramming is a potential therapeutic approach for managing IPF.

## Introduction

Idiopathic Pulmonary Fibrosis (IPF) is a progressive, aging-related, lung disease characterized by excess matrix deposition, lymphangiogenesis, macrophage accumulation, and epithelial cell remodeling^1^. Genetic and environmental factors like smoking, air pollution, or chitin accumulation^2^ interact to drive IPF lung remodeling and fibrosis. Mutations in telomere associated genes are found in a subset of familial and sporadic IPF patients^3–7^. Variants or mutations in epithelial cell-associated genes (Muc5B, Abca3, Sftpa2, Dsp) also increase IPF risk. Short telomeres in alveolar type 2 (AT2) cells are found in all IPF patients^8–11^. While telomere dysfunction is considered a molecular driver of lung remodeling and fibrosis, the downstream molecular mechanisms causing tissue remodeling and fibrosis remain poorly understood.

Telomeres are repetitive nucleotide structures at chromosome ends. The telomere shelterin complex protein Telomere Repeat Binding Factor 1 (TRF1 / Terf1) binds to TTAGGG repeats and is critical for maintaining the protective shelterin cap^12^. Conditional deletion of TRF1 in AT2 cells is sufficient to cause elements of remodeling and fibrosis found in IPF lung^13,14^. Because AT2 cell telomere dysfunction is implicated as an initiator of IPF, mice lacking AT2 cell TRF1 may help advance understanding of the molecular mediators that initiate lung remodeling and fibrosis in IPF.

Single cell RNA-sequencing (scRNA-seq) studies have identified novel epithelial cell subtypes in IPF such as intermediate Krt8+^15^, pre-alveolar transitional cell state (PATS)^16^ and Krt5-/Krt17+^17^ matrix-producing cells. Why these epithelial cell subtypes emerge and whether telomere dysfunction contributes to their accumulation or initiation of epithelial cell remodeling leading to IPF is not known. Conceptually, the pathogenesis of IPF transitions through multiple stages like genetic risk, initiation and progression with various pathologic cell types and mediators dominating each stage^18^. In the initiation stage epithelial cell failure due to telomere dysfunction, and senescence reprogramming appear to dominate. In response to telomere dysfunction, activation of a DNA damage response (DDR) is a proximal event that is sensed by ATM, ATR and DNA-PK kinases leading to p53 activation and cell cycle arrest^19,20^. This study aims to understand the early molecular mediators of AT2 cell telomere dysfunction that initiate lung remodeling and fibrosis and elements of remodeling attributable to AT2 cell telomere dysfunction. The findings in this manuscript suggest the DNA damage response to AT2 cell telomere dysfunction initiates p53-dependent progressive lung remodeling and fibrosis in IPF lungs.

## Results

### Single cell analysis of lung epithelial cells defines epithelial cell remodeling in response to AT2 cell telomere dysfunction

To understand how AT2 cell telomere dysfunction contributes to lung epithelial cell remodeling, lungs were harvested from SPC-creTRF1^flox/flox^ mice treated with tamoxifen for 2 months (early time-point; no fibrosis) or 8 months (late time-point; established fibrosis). Epcam+ cells were sorted for single cell RNA sequencing (scRNA-seq) (Figure 1A). The Uniform Manifold Approximation and Projection (UMAP) of epithelial cell clusters from scRNA-seq data revealed epithelial cell heterogeneity between control, early, and late time-points (Figures 1B and S1A) identifying 27 clusters (Figure. S1B) based on cluster-specific makers (Figure S2). Control lungs yielded the expected epithelial cell populations including AT2 cells which separated into two subtypes based on Lyz1 expression. In addition, an intermediate AT2 cell cluster (Cluster 15) connected control AT2 sub-population with the Lyz1 high AT2 cell cluster (Figure 1C).

**Figure 1.**
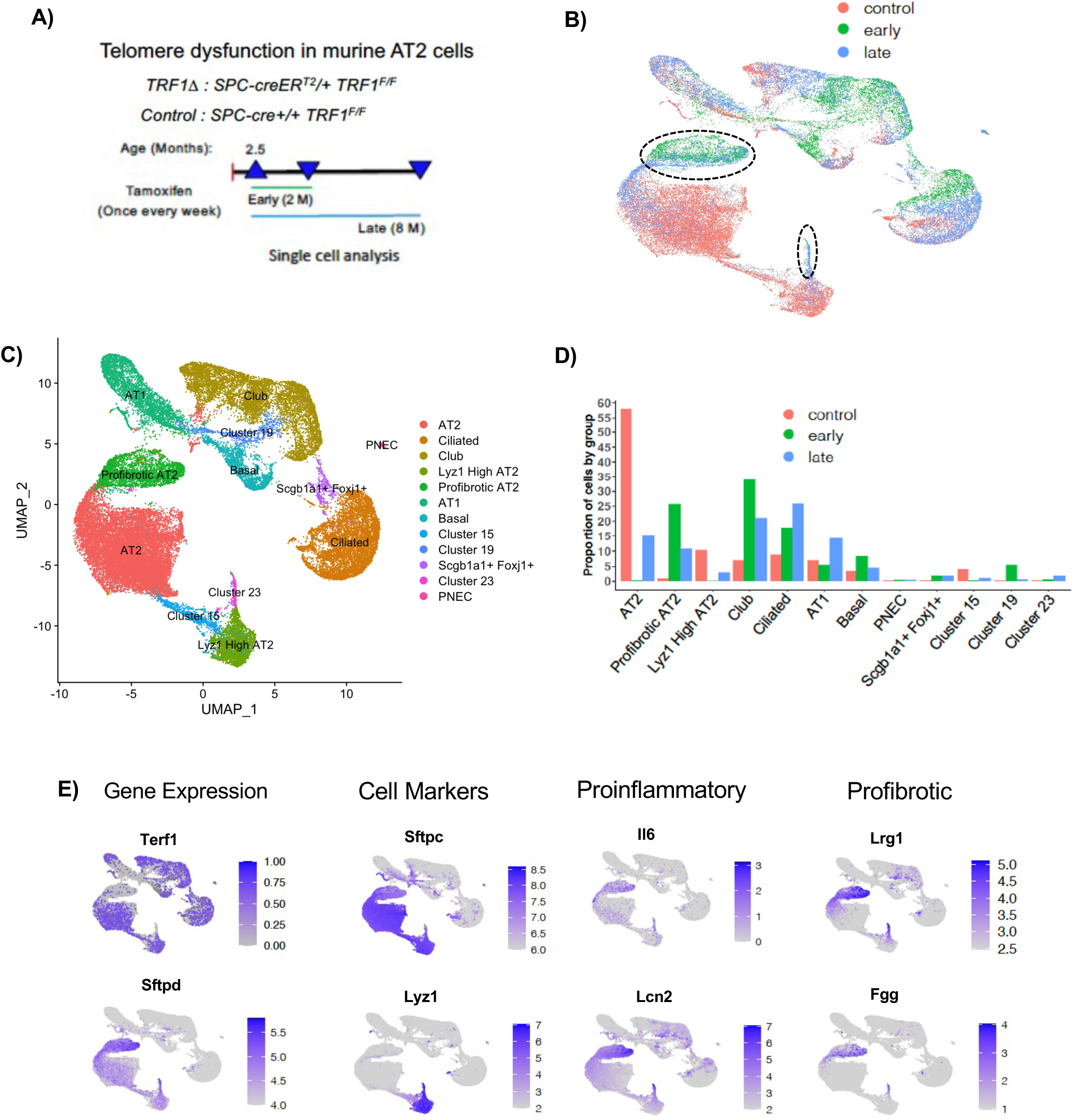
Single cell distribution of lung epithelial cells after telomere dysfunction in AT2 cells. A) Schematic of telomere dysfunction model in SPC-creERT2 mice to conditionally delete TRF1. Tamoxifen was injected at 250 mg/kg body weight. Lungs were harvested from SPC-creTRF1F/F mice for single cell-RNA sequencing of epithelial cells at Early timepoint (2 months) (n=1) or Late (8 months) timepoint (n=2) and age-matched TRF1F/F controls (n=2). B) UMAP plot showing lung epithelial single-cell distribution comparing control with early (2M), late (8M) phenotypes. C) UMAP plot showing cellular identity based on cluster-specific markers and clusters of unknown identity. D) Bar graph showing cell proportions in each cell cluster comparing groups. E) Feature plots displaying expression level of Terf1, Lyz1, AT2 cell-specific markers, pro-inflammatory and pro-fibrotic markers within lung epithelial cells. Positive gene expression (blue color gradient) or no expression (grey). Color scale indicates fold expression.

In SPC-creTRF1^flox/flox^ mice subjected to AT2 cell telomere dysfunction following tamoxifen treatment, novel AT2 cell populations emerged at both early and late time-points (Figure 1B, marked by dotted lines), identified as profibrotic AT2 and a second cluster of pathologic AT2 cells in cluster 23 (Figure 1C). The profibrotic AT2 cell cluster and cluster 23 lacked TRF1 (Terf1) expression confirming TRF1 deletion from pathologic AT2 cells (Figure 1E). The absence of TRF1 in pathologic AT2 cells was associated with elevated surfactant protein D (Sftpd), pro-inflammatory molecules like Lipocalin2 (Lcn2) and Interleukin-6 (Il-6), pro-fibrotic molecules like leucine-rich alpha-2-glycoprotein-1 (Lrg1), and Fibrinogen gamma chain (Fgg) (Figure 1E). Quantification of cell proportions showed a marked decrease in normal AT2 cells following telomere dysfunction at early and late time-points. The proportions of club and ciliated cells increased at both early and late time-points while basal cell proportions only increased slightly at the early time-point (Figure 1D). This demonstrates that epithelial cell remodeling following AT2 cell telomere dysfunction is a dynamic process, with different epithelial populations expanding over time to compensate for dysfunctional AT2 cells.

### Airway epithelial cell response to AT2 cell telomere dysfunction

To visualize club cells on the UMAP, a feature plot was generated for Scgb1a1 (Figure 2A). Scgb1a1 expression was widespread in early and late lungs compared to control indicating club cell expansion. A subset of club cells expressing Scgb3a1 appeared in lungs with telomere dysfunction with little expression in control. (Figure 2A). Immunostaining lung tissues with Scgb1a1 and Scgb3a1 antibodies demonstrated that Scgb1a1 expressing club cells were located in the alveolar parenchyma of tamoxifen treated SPC-creTRF1^flox/flox^ mice but not in control mice (Figure 2B, upper panels). Control mouse lungs had few Scgb3a1-expressing club cells in larger airways and none in smaller airways (Figure 2B, lower panels). In contrast, there was an expansion of Scgb3a1-expressing club cells in both large and small airways of tamoxifen treated SPC-creTRF1^flox/flox^ mice, suggesting that two club cell populations expand in response to AT2 cell telomere dysfunction, possibly to repopulate lung regions with dysfunctional AT2 cells. To determine if club cell expansion is similarly present in IPF lungs, normal and IPF lungs were immunostained for Scgb1a1. Immunofluorescent staining showed greater proportions of Scgb1a1 immunoreactive club cells in IPF airways and adjacent lung parenchyma (Figure 2C). In SPC-creTRF1^flox/flox^ mice, genes such as Tff2, Muc5b, Gp2, Dmbt1, Tafa4 and Agr2 localized to the Scgb3a1 cells. These genes are known to have functional role in proliferation, migration, differentiation and defense mechanisms (Figure. 2D). To test for differences between club cells from control and those that expand in response to AT2 cell telomere dysfunction, differential expression analysis was performed. Top ten upregulated and down-regulated genes are displayed (Figure 2E). To signify the importance of all differentially expressed genes in club cells, KEGG pathway analysis indicated enriched pathways (Figure 2F).

**Figure 2.**
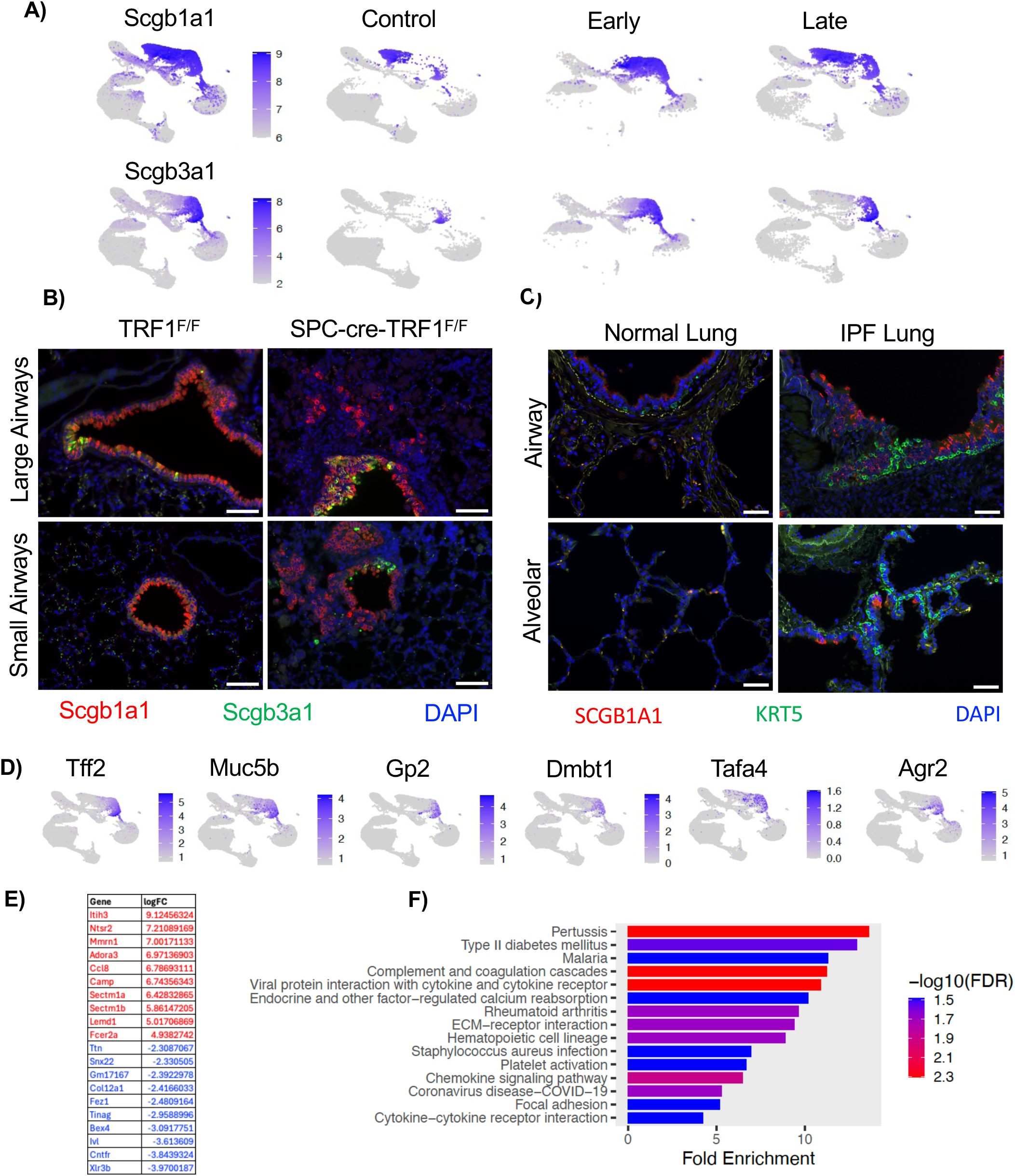
Airway epithelial cell response to telomere dysfunction in AT2 cells. A) Feature plots showing expression of Scgb1a1 and Scgb3a1 (Left panel: merged; Right 3 panels: split by group). Color scale indicates fold expression. Blue=positive expression; Grey = No expression. B) Immunofluorescence staining of Scgb1a1 (red) and Scgb3a1 (green) in TRF1F/F control, SPC-creTRF1F/F (8 Months) after telomere dysfunction. Upper panels are large airways, and lower panels are small airways. Scale=200𝜇m. C) Immunofluorescence staining of human normal and IPF lungs with KRT5 (red) and SCGB1A1 (green). Scale=25𝜇m. D) Feature plots representing gene expression in the Scgb3a1 positive population. E) Top 10 differentially expressed genes (up-regulated: red and down-regulated: blue) comparing club cells from both groups (control versus those with telomere dysfunction in AT2 cells). F) KEGG pathway analysis of differentially expressed genes shows dysregulated pathways in club cell cluster.

### Telomere dysfunction causes broad reprogramming of AT2 cells

Gene expression was compared in AT2 cells from control, early and late populations. The analysis demonstrated that both profibrotic and cluster 23 AT2 cells (comprising of early and late populations) exhibited higher expression levels of DNA damage, (Ccnd1, Ccng1, Trp53inp2), senescence (H2-Q7, Slpi, Itih4, Pigr, Mapk13) and lung remodeling (Retnla, Lrg1, Orm1, Serpine2, Timp2, Lbp, Cd44, Fgg) genes in response to telomere dysfunction (Figure 3A).

**Figure 3.**
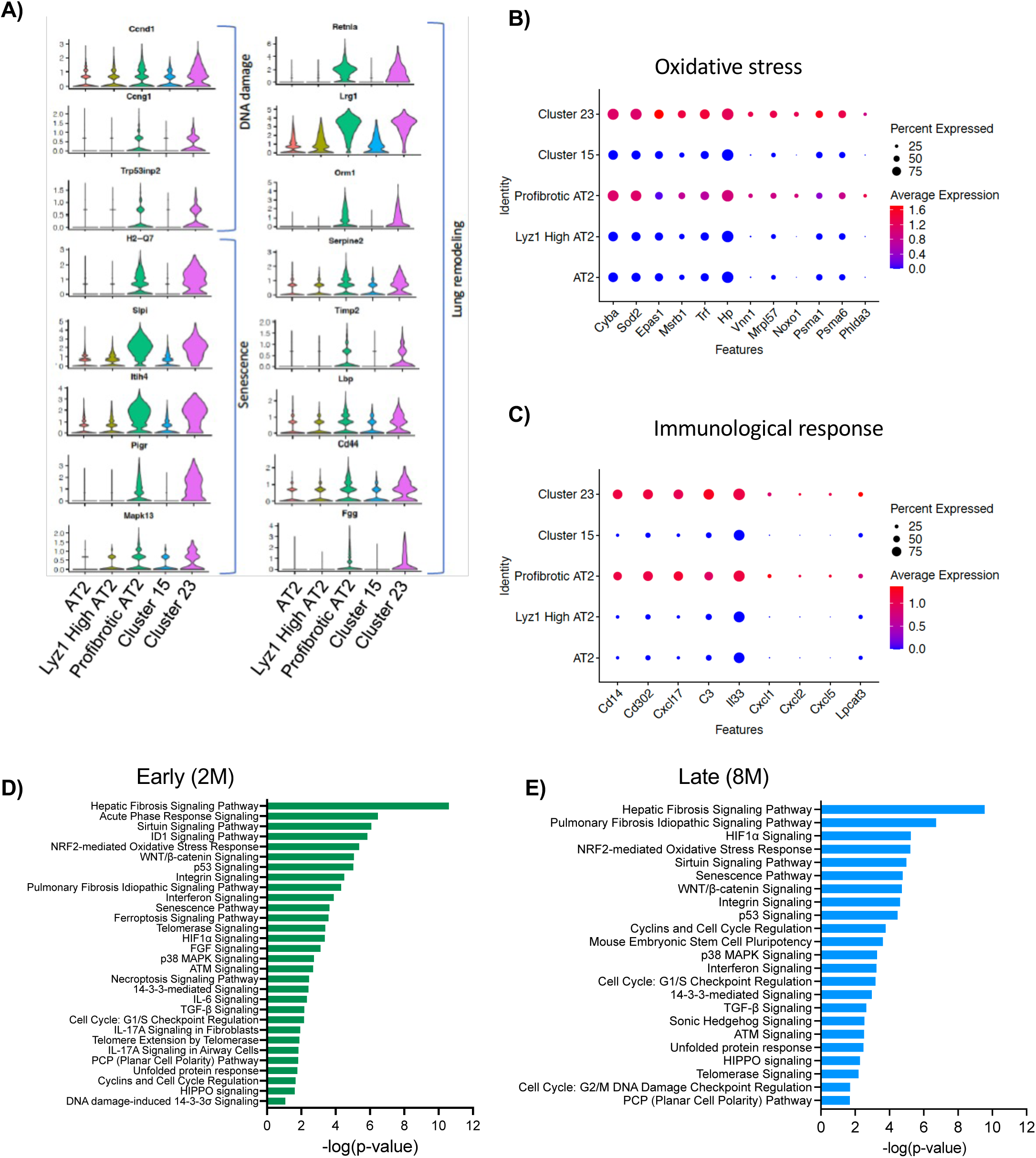
Gene expression changes and canonical signaling pathways in AT2 cells after telomere dysfunction. A) Violin plots displaying expression levels of DNA damage, senescence and lung remodelingmarkers in AT2 cells 8 months after telomere dysfunction. Dot plots showing markers of B) oxidative stress and C) Immunological response. Color legend displaying blue (lowest) to red (highest) expression gradient. Size of the dot indicates percent cells expressing the gene. D and E) Ingenuity pathway Analysis (IPA) showing top dysregulated pathways at D) Early (2 months) and E) Late time point (8 months) after telomere dysfunction.

Cellular stress, including oxidative stress, is a characteristic of IPF AT2 cells. In the SPC-cre TRF1^flox/flox^ mice with telomere dysfunction, there was upregulation of oxidative stress response genes such as Sod2, Noxo1 and Cyba in profibrotic AT2 cells and cluster 23 AT2 (Figure 3B). In addition, these AT2 cells demonstrated increased expression of chemokines (e.g. Cxcl1, Cxcl2), cytokines (e.g. IL-6) and immunomodulators like Cd302 and Cd14, all known to attract leukocytes (Figures 3C and S3A and S3B). These findings suggest that AT2 cell telomere dysfunction, DNA damage response, and senescence reprogramming may be contributing to oxidative stress and production of proinflammatory mediators in IPF AT2 cells. Differentially expressed genes were analyzed using Ingenuity Pathway analysis (IPA) to identify dysregulated signaling pathways attributable to telomere dysfunction. The top dysregulated signaling pathways in early and late AT2 cells included hepatic and pulmonary fibrosis pathways, p53, Wnt, Tgf-β, hippo and cell cycle pathways^21,22^. These pathways have been shown to be dysregulated in IPF AT2 cells, suggesting their dysregulation may be a consequence of telomere dysfunction (Figures 3D and 3E).

### Trajectory analysis of epithelial cells defines transition from quiescent to pathologic epithelial cell states

Trajectory analysis using slingshot^23^ revealed the directionality of AT2 cell phenotypic change from cluster 0 (control AT2) through cluster 1, ending in profibrotic AT2 cell clusters 6 and 10 (Figure 4A) following onset of AT2 cell telomere dysfunction. Gene expression changes along the trajectory highlight the loss of AT2 cell markers and acquisition of airway epithelial cell markers (e.g. Scgb1a1, Scgb3a1) with increased Sftpd expression (Figure 4B). In addition, profibrotic AT2 cells show increased expression of potential senescence associated secretory phenotype (SASP) genes (Mapk13, Il6, Serpine2, Cxcl2, Cxcl5) and a pro-fibrotic gene signature (Lrg1, Dmkn, Slpi, Fgg). The analysis also demonstrated a decrease in AT2 cell progenitor genes (Fzd5, Fzd8, Sox4) and cell cycle maintenance genes (Bex2, Bex4) in profibrotic AT2 cells, suggesting a failure of genes regulating AT2 cell self-renewal. The trajectory from normal Lyz1 high AT2 cells to cluster 23 AT2 cells show overlap with AT2 cell trajectory with the expression of Lyz1 being the primary difference (Figures 4C and 4D).

**Figure 4.**
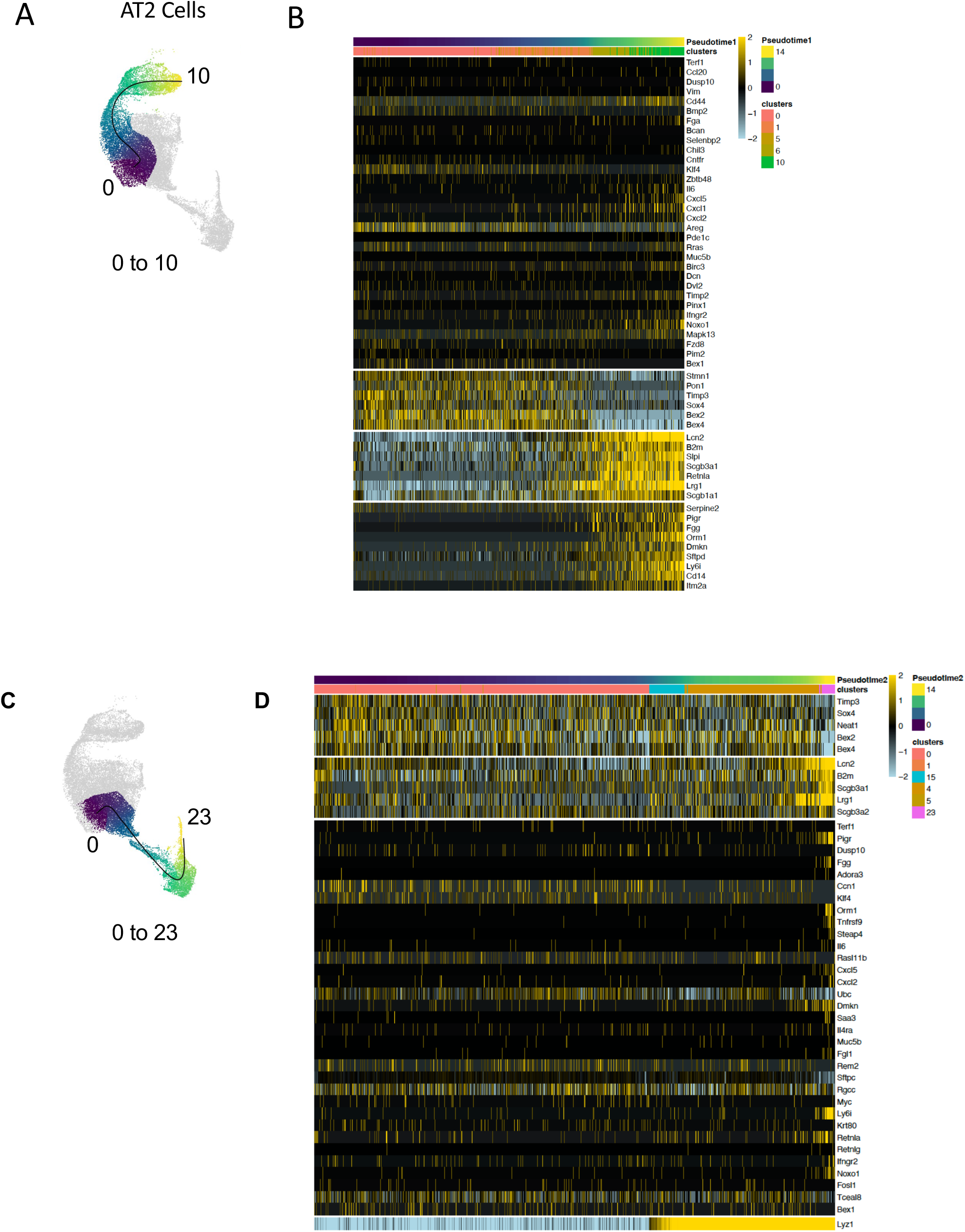
Trajectory of epithelial cells displaying the path from control AT2 cells to profibrotic AT2 cells with telomere dysfunction. A) Slingshot trajectory showing the path of control AT2 cells (purple) towards dysfunctional AT2 cells (yellow) (Pseudotime 1). B) Heatmap showing differentially expressed genes along the trajectory from cluster 0 to clusters 6, 10. Color scale indicating relative expression from −2 (blue) to 2 (yellow). C) Slingshot trajectory showing the path of control AT2 cells (purple) to Lyz1 high AT2 cells (yellow) (Pseudotime 2). D) Heatmap showing differentially expressed genes along the trajectory from cluster 0 to 23. Color scale indicating relative expression from −2 (blue) to 2 (yellow).

Trajectory analysis of club cells identified gradual transition from control club cells expressing Scgb1a1 to a pathologic club cell subset expressing Scgb3a1 that expressed markers of proliferation (Figures S4A and S4B). This Scgb3a1 subset expressed higher levels of defensins (Bpifa1, Bpifb1), mucins (Muc5b), keratins (Krt19) and regenerating family member (Reg3g). The trajectory leading from club cells to basal cells has gene signature including Krt5, Krt17, Trp63, Phlda3, Mir205hg, Wnt4 (Figures S4C and S4D). This trajectory has cluster 19 intermediate. Cluster 19 has features of proliferating epithelial cells with markers such as Clu, S100A6, Krt19, Krtap17-1, Cldn4, Fxyd3 and was predominantly found at early time-point (Figure S5).

### Telomere dysfunction-induced gene expression changes in AT2 cells over time

Differential expression analysis of AT2 cell clusters identified gene expression variations at early and late time-points. Some genes were enriched only at early time-point (Figure 5A) and disappeared by the late time-point. Early-specific genes included those involved in necroptosis (Ripk3), cell cycle (Gadd45g, Ccnb1), extracellular matrix (ECM) receptor interaction (Cd44, Itgb3, Fn1, Lamc2, Itgb6), glycosaminoglycan biosynthesis (Dse), cytokine-cytokine receptor interactions (Gdf11, Ccl5, Tgfb2, Tnfrsf11b) and epithelial cell differentiation (Wnt4). Genes with decreased expression at the early time-point include Fzd5, Fzd8, Hes2 and Bmp3 (Figure 5A). KEGG pathway analysis of early-specific genes indicated enrichment of pathways involved in ECM-receptor interaction, cytosolic DNA sensing, cell cycle, cell differentiation among others (Figure 5B). Conversely, there was a late-specific gene signature in AT2 cells. Late-specific genes include those involved in activation of cell contraction (Abra), cell adhesion (Cdh5), matrix metalloproteinases (Mmp3), activation of TGF-beta signaling (Acvrl1 or Alk1), or epithelial to mesenchymal transition (Vim, Inmt) (Figure 5C). KEGG pathway analysis of late-specific genes indicated enrichment of pathways involved in protein digestion and absorption, ECM-receptor interaction, and PI3K-Akt signaling pathways (Figure 5D).

**Figure 5.**
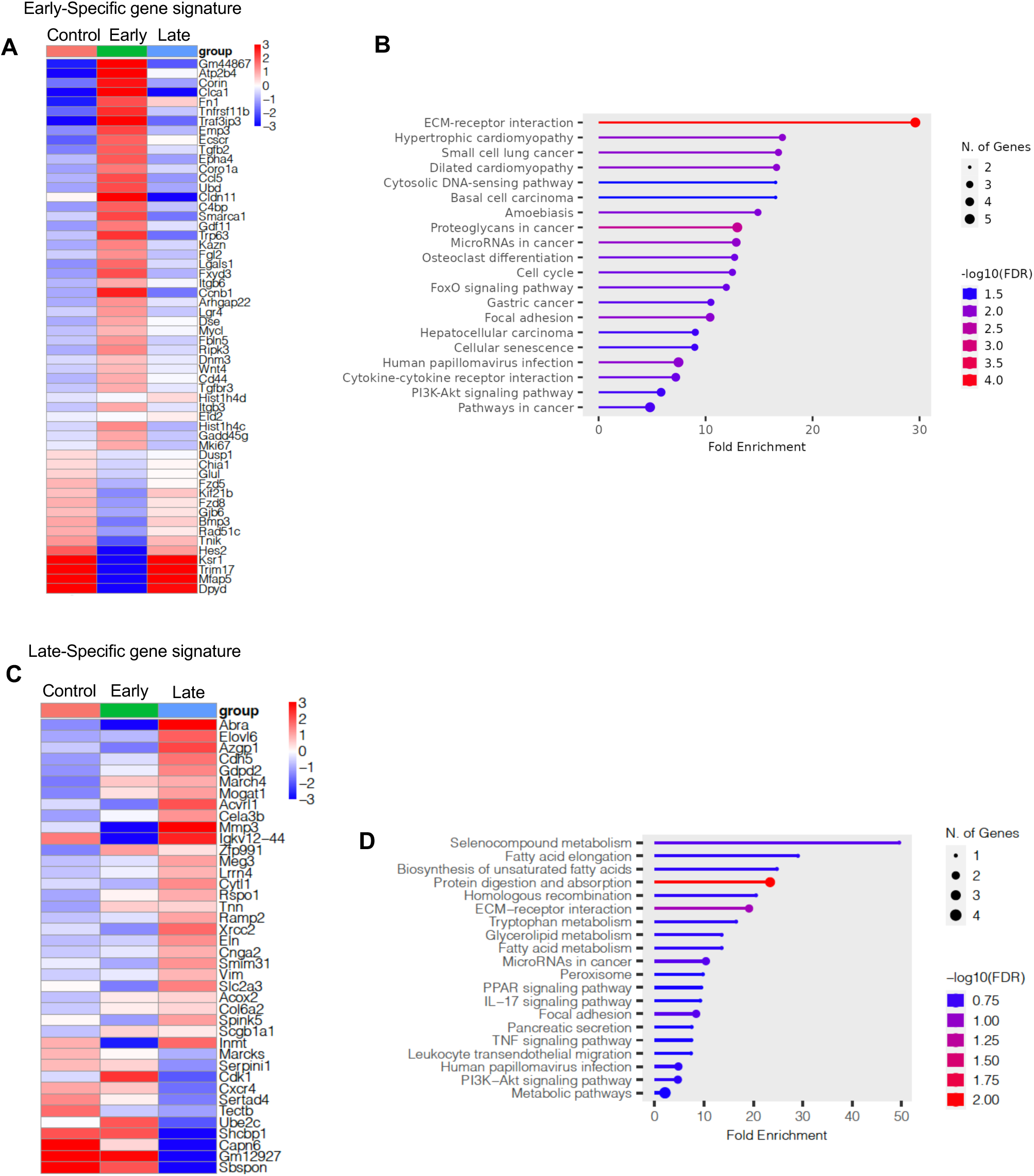
Gene expression changes and canonical signaling pathways in AT2 cells after telomere dysfunction. A) Heat map reflecting differentially expressed genes comparing control, early and late groups and are exclusive to early phenotype showing upregulated (red) and downregulated (blue) genes. B) KEGG pathway analysis of early-specific differentially expressed genes depicting fold enrichment. C) Heatmap showing gene expression changes comparing control, early and late groups and are exclusive to late phenotype. Highest expression (red) and lowest in (blue). D) KEGG pathway analysis of differentially expressed genes in late-specific timepoint. Size of the dot indicates number of genes in the dataset that overlap with the genes in the pathway. Color scale indicates enrichment FDR.

Expression of some genes peaked at the early time-point then were expressed at lower levels later. Early-peaking genes included proinflammatory (Retnla, Orm3, Saa3, Tnf, Tnfrsf9, Lcn2), profibrotic (Lrg1) and telomere maintenance (Zbtb48) genes (Figure S6A). KEGG pathway analysis of the upregulated genes showed IL-17 pathway as top enriched pathway with genes including cytokines (TNF, Il6), chemokines (Cxcl5, Cxcl1), Muc5b, and Lcn2. Other enriched pathways include cytokine-cytokine receptor interactions (Ifngr2, Tnfrsf9, Cxcl17) and cellular senescence (Il6, Mapk13) (Figure S6B). Genes suppressed specifically at early stage involved cancer signaling Rap1, Ras pathways (Map2k3, Fgfr1, Fgfr3). mTOR pathway inhibitor Deptor, p53 pathway inhibitor Mdm1 are suppressed promoting activation of mTOR and p53 pathways respectively (Figures 6C and 6D). Finally, certain gene changes are in a continuum from early to late stage. The notable gene changes include upregulation of Ccl2, Hif3a and downregulation of Terf1 expression (Figure S7A). The GO biological processes involve positive regulation of PDGF receptor signaling (F7, Hip1), telomeric loop disassembly (Terf1), myeloid leukocyte migration (Pde4b, Ccl2) (Figures S7A and S7B). Collectively, these data show that AT2 cells continue to change gene expression over time following onset of telomere dysfunction.

**Figure 6.**
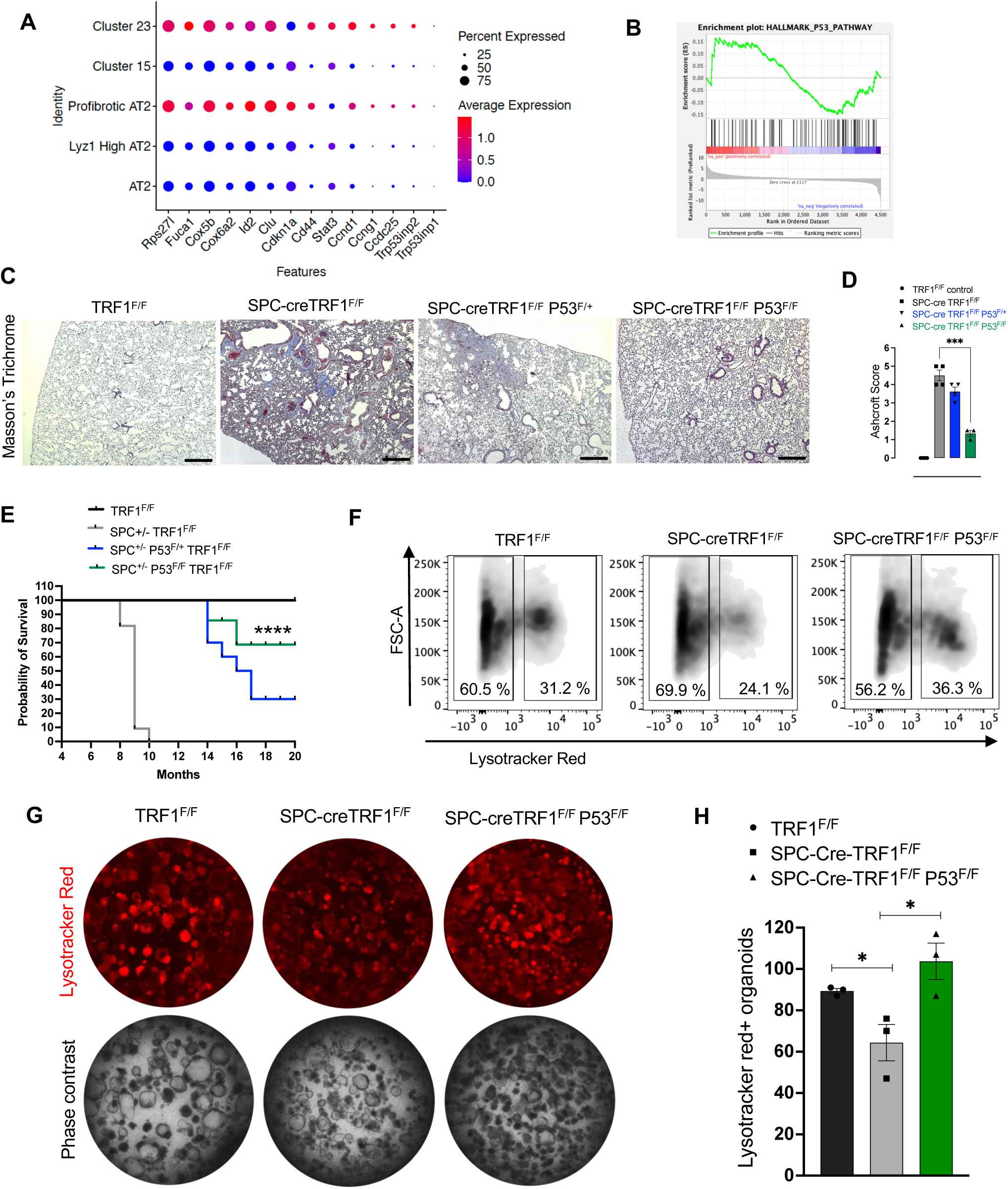
Ablation of P53 in AT2 cells ameliorated lung fibrosis and revived organoid growth. A) Dot plot showing p53-effector gene expression in mouse model of telomere dysfunction. Color panel indicates expression level and size of the dot indicates percent cells expressing the gene. B) GSEA analysis of differentially expressed genes from scRNA-seq data of AT2 cell clusters showing p53 pathway enrichment. C) Masson’s trichrome staining showing collagen deposition (blue). Scale bar = 800𝜇m. D) Quantification of lung remodeling and fibrosis by Ashcroft method. n=3-4 mice per group; p-value<0.001***. E) Kaplan Meier survival curve indicating the probability of survival. N=7-10 mice per group; ****P<0.0001 (log-rank test) comparing SPC-creTRF1F/F and SPC-creTRF1F/F P53F/F mice. F) Flow cytometry analysis of lysotracker red positive cells. G) Organoids derived from co-culture of NIH-3T3 fibroblasts with lung epithelial cells from TRF1F/F control, SPC-creTRF1F/F or SPC-creTRF1F/F P53F/F mice after telomere dysfunction. Representative images captured on day 14. H) Quantification of lysotracker red positive organoids n=3 mice per group; pvalue<0.05*.

### Ablation of p53 in SPC expressing cells ameliorates fibrosis, improves survival, and promotes organoid growth

Targets of a p53 gene signature^24^ and GSEA analysis of differentially expressed genes representing a p53-activation score were upregulated in profibrotic and cluster 23 AT2 cells indicating activation of the p53 pathway and its target genes (Figures 6A and 6B). To test whether p53 activation drives lung remodeling and fibrosis in SPC-creTRF1^flox/flox^ mice, they were crossed to p53^flox^ mice to genetically delete p53 in SPC-expressing cells. Consistent with prior reports^13^, SPC-creTRF1^flox/flox^ mice treated with tamoxifen developed lung remodeling and fibrosis spontaneously. In contrast, telomere dysfunction in SPC-creTRF1^flox/flox^p53 ^flox/wt^ mice exhibited less lung remodeling and improved survival whereas complete ablation of p53 in SPC expressing cells (SPC-cre TRF1^flox/flox^ p53 ^flox/flox^ mice) hereafter referred as p53 conditional knockout (p53-CKO) did not exhibit lung remodeling and fibrosis as shown by decreased masson’s trichrome staining, lower Ashcroft score, and improved survival (Figures 6C, 6D and 6E). To examine the effect of p53 ablation on AT2 cell burden, lung epithelial cells harvested from TRF1^flox/flox^, SPC-creTRF1^flox/flox^ and p53CKO mice were analyzed by flow cytometry. While SPC-creTRF1^flox/flox^ mice showed a decrease in lysotracker red positive cells compared to TRF1 controls, p53CKO mice demonstrated an increase in lysotracker red positive cells suggesting that absence of p53 in SPC-expressing cells maintained AT2 cells in p53CKO mice (Figure 6F). Epcam positive cells from TRF1^flox/flox^, SPC-creTRF1^flox/flox^ and p53-CKO mice were plated on matrigel along with NIH3T3 fibroblasts to form lung organoids. After 14 days in culture, SPC-creTRF1^flox/flox^ epithelial cells yielded fewer lysotracker positive organoids. In contrast, p53-CKO mice had significantly more lysotracker positive organoids (Figures 6G and 6H).

### p53 inhibition enhances organoid formation and limits profibrotic gene expression in IPF AT2 cells

p53 is a DNA damage signaling mediator activated in response to telomere dysfunction^25^. To investigate the role of p53 in IPF AT2 cell transdifferentiation and failure, lysotracker red positive cells were collected from IPF lung and cocultured with MRC5 fibroblasts for 14 days. (Figure 7A) with either DMSO or the p53 inhibitor (p53i) pifithrin alpha hydrobromide at 5μM. IPF AT2 cells cultured in the presence of Pifithrin-α hydrobromide showed increased colony forming efficiency of lysotracker+ organoids (Figures 7B and 7C) with no change in total colonies (Figure 7D). Single cell analysis of day 14 organoids shows that p53 inhibition preserved the AT2 cell identity (Figure 7E), decreased expression of DNA damage response and senescence markers (Figure 7F) in epithelial cells and decreased TGFb signaling (Figure 7G), and pathologic collagen expression in fibroblasts (Figure 7H).

**Figure 7.**
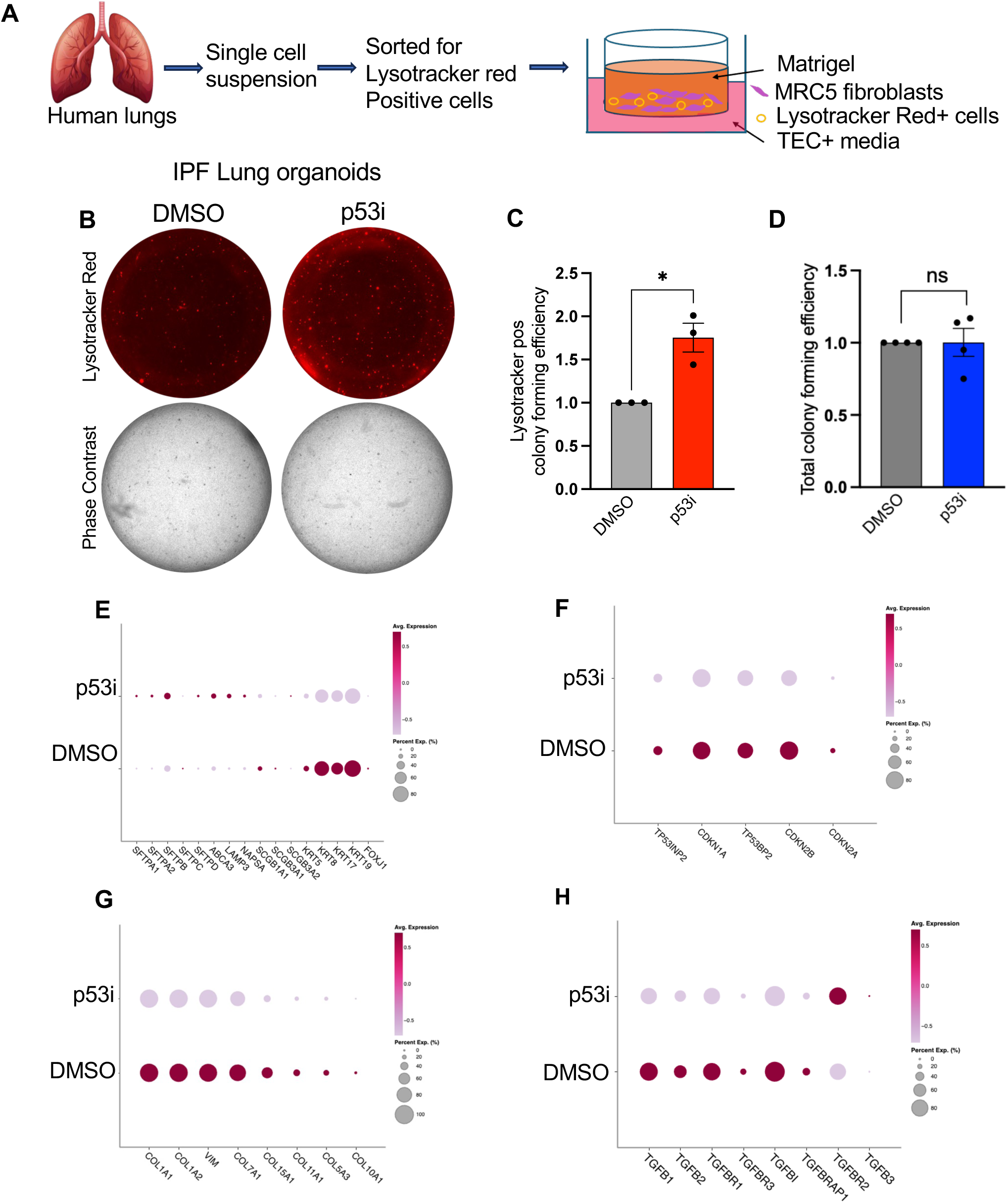
Effect of p53 inhibition on IPF lungs in a three-dimensional (3D) lung organoid system and single cell sequencing. **A)** Schematic depicting the human 3D lung organoid culture **B)** Lysotracker positive epithelial cells sorted from IPF lung were co-cultured with MRC5 fibroblasts. Organoids were treated with DMSO or 5𝜇M P53 inhibitor (p53i) Pifithrin-alpha hydrobromide. Organoids were cultured for 14 days. Lysotracker red (top panel) positive colonies and phase contrast (bottom panel) colonies shown are representative of day 14. **C,D)** Quantification of **C)** lysotracker red positive and **D)** total colony forming efficiency. Ns=not significant, *p<0.05, n=3. **E-H)** Dot plots showing differential expression between p53i and DMSO for **E)** epithelial cell markers **F)** DNA damage response and senescence in epithelial cells **G)** Collagen expression in fibroblasts. **H)** TGF beta pathway gene expression in fibroblasts. Color scale indicates fold expression. Brown=positive expression; Light pink = No expression. Size of the dot indicates percent cells expressing the gene.

## Discussion

IPF is a complex disease involving various pathologic processes including short telomeres and senescence reprogramming of AT2 cells, alveolar epithelial cell remodeling, expression of profibrotic molecules, immune cell infiltration and appearance of an activated profibrotic (CTHRC1) fibroblast population^26^ leading to excess lung matrix and collagen deposition^1^. The chronology of these events and the earliest molecular changes in IPF are unresolved limiting the development of therapies targeting molecular pathways that initiate the pathologic process.

Unfolding the early molecular events in IPF is essential for understanding the pathologic continuum of IPF. Most studies concern fibroblast activation, a late event in the disease^18^. Genetic studies^27^ and the observation of short telomeres in AT2 cells of IPF patients have led to investigations of the role of AT2 cell failure in IPF^8^. Lung fibrosis models commonly use bleomycin to injure the lungs. However, this model has limitations: lung injury precedes fibrosis, which resolves unlike IPF, which is progressive. To address these limitations, we developed a mouse model of progressive lung fibrosis initiated by telomere dysfunction in AT2 cells (SPC-creTRF1^flox/flox^ mice)^13^. To understand the cellular and molecular events, single-cell RNA sequencing was performed at early and late time-points after initiating telomere dysfunction in AT2 cells of SPC-creTRF1^flox/flox^ mice. The emergence of a profibrotic AT2 cell signature at an early time-point, when no histopathological fibrosis is present, suggests that profibrotic reprogramming of AT2 cells is the earliest molecular change in IPF pathogenesis.

AT2 cells have been reported to undergo reprogramming to a Krt8+ transitional stem cell state^15^, DATPs^28^ or Krt5 expressing basal cells^29^ in response to various stressors^30^. Although these pathologic epithelial cell subtypes are present in IPF lungs, the molecular drivers of the reprogramming are unresolved. In this model, AT2 cells with telomere dysfunction transitioned to a profibrotic AT2 cell phenotype that expressed markers of DNA damage, senescence, oxidative stress and lung remodeling. Many of these markers are expressed in IPF AT2 cells^10,31,32^ and pathologic epithelial cells, suggesting that telomere dysfunction is a molecular driver that initiates AT2 cell dysfunction and disrupts epithelial cell homeostasis.

In response to AT2 cell telomere dysfunction, there is compensatory expansion of bronchial epithelial cells, including Scgb1a1 and Scgb3a1-expressing club cells and ciliated cells^33^. In SPC-creTRF1^flox/flox^ mice, club cells are the first airway associated epithelial cell to appear in the alveolar compartment. This suggests that club cells may be early responders to the failure of IPF AT2 cells due to telomere dysfunction and senescence reprogramming. Unlike bleomycin-treated lungs or explanted IPF lungs^29^, Krt5+ basal cells were not present in the remodeled alveolar parenchyma of SPC-creTRF1^flox/flox^ mice. This absence suggests that additional external stressors may be required to cause club cell failure, enabling basal cells to emerge as second-line responders to AT2 cell failure. In IPF patients, exposure to environmental stressors such as tobacco smoke or pollutants could result in loss of club cells, enabling basal cells to expand.

Pathologic features of IPF include telomere dysfunction, activation of a DNA damage response (DDR) and senescence reprogramming of AT2 cells^31,34^ Sensors of telomere dysfunction include ATM, ATR and DNA-PK^19,20^. These DNA damage response kinases signal the presence of telomere dysfunction, inhibit the cell cycle, and promote senescence reprogramming in part through p53. In this study, we found that genetic deletion or inhibition of p53 promoted a “healthier” AT2 cell phenotype, characterized by higher expression of AT2 cell markers and lower expression of basal cell markers in AT2 cells from SPC-creTRF1^flox/flox^ mice and IPF patients. Interestingly, organoid experiments did not yield higher numbers of organoids, suggesting that p53 inhibition had little effect on cell division. These data suggest that the p53-dependent component of AT2 cell senescence reprogramming response to telomere dysfunction is reversible and inhibition of pathologic p53 signaling promotes the health of senescent AT2 cells^35^.

Genetic deletion of p53 in SPC expressing cells with telomere dysfunction prevented lung fibrosis in SPC-creTRF1^flox/flox^ mice. This appears to be in part due to shifting of the senescent AT2 cells to a more normal phenotype. In addition, IPF organoid experiments demonstrated less matrix and collagen production by fibroblasts co-cultured with p53 inhibitors. These findings suggest that senescent AT2 cells directly activate fibroblasts and initiate excess matrix production and that this activation may in part be due to TGF beta because TGF beta signaling mediators were also expressed at a lower levels in p53-inhibited AT2 cell organoid fibroblasts. Whether this crosstalk continues to contribute to fibrosis progression at later disease stages or in the context of lung injury remains to be investigated.

In normal physiology, p53 regulates cellular metabolism and maintains cell homeostasis^36^. Within normal lung, p53 regulates airway epithelial cell progenitor behavior and differentiation of AT2 cells to AT1 cells^16,37^. In pathologic conditions, p53 signaling is activated in response to stressors such as DNA damage, heat shock, hypoxia, or telomere dysfunction among others. In response to telomere dysfunction, p53 levels are durably increased and associated with cell cycle arrest and senescence reprogramming^38^. Senescence of AT2 cells mediated by pathologic upregulation of p53 has been implicated in lung fibrosis^39,40^. These findings and our data support the idea that targeting pathologic p53 may reprogram dysfunctional AT2 cells and improve cellular homeostasis.

There is increasing interest in targeting senescent epithelial cells as drivers of pulmonary fibrosis, either by eliminating them using senolytics or reversing the senescence phenotype using senomorphics^41^. The data presented in this manuscript suggest that inhibiting enhanced pathologic p53 activity may favorably reprogram senescent AT2 cells driven by telomere dysfunction, suggesting this could be considered a therapeutic approach for limiting an early profibrotic event in IPF. Although a common concern is that such a therapy may promote cancer, inhibition of p53 alone is not sufficient to cause cancer^42^ as additional gene mutations are required. Whether p53 inhibition can be done globally or via cell-specific inhibition to limit risk of deleterious effects of physiologic p53 inhibition requires further study.

## Resource Availability

### Lead Contact

Further information and request for resources and reagents should be directed to the lead contact, Paul Wolters

### Materials availability

This study did not generate any new unique reagents

### Code availability

The code of the analysis is on https://github.com/gladstone-institutes/GB-RN-1264/tree/main

## Methods

### Mice

SPC-Cre/ERT2,rtTA mice, were obtained from Hal Chapman^43^. p53^LoxP^ mice were purchased from Jackson laboratory. All the mice are on C57BL/6 background. Mice were bred and housed in pathogen-free conditions in accordance with the guidelines of Laboratory Animal Resource Center (LARC). All animal procedures were carried out using protocols approved by the Institutional Animal Care and Use Committee at the University of California, San Francisco.

### Tamoxifen administration

Tamoxifen (Toronto Research Chemicals, Cat # T006000), 250 mg / kg body weight was injected once per week by intraperitoneal route to *SPC-creTRF1^F/F^* mice, *SPC-creTRF1^F/F^*p53^F/F^ mice and *TRF1^F/F^* littermate controls beginning at 10 weeks age.

### Antibodies and Reagents

Commercially available antibodies were purchased from following vendors. Rabbit polyclonal SPC antibody (Abcam; Cat # ab3786), rat monoclonal human Scgb1a1 (Cat # MAB4218), mouse monoclonal human Scgb3a1 (Cat # MAB2790), goat polyclonal Scgb3a1 (AF2954) from R&D Biosystems, rabbit polyclonal CC10 (sc-25555), Keratin 5 chicken polyclonal antibody (Biolegend, Cat # 905901), cell viability dye sytox blue (Thermofisher, Cat # S34857). Fluorescent secondary antibodies donkey anti-chicken Alexa fluor 488 (Jackson immuno research, Cat # 703-545-155), donkey anti-mouse Alexa Fluor 488 (Life Technologies, Cat # A21202), donkey anti-goat Alexa Fluor 546 (Life Technologies, Cat # A11056), donkey anti-rat Alexa Fluor 594 (Life Technologies, Cat # A21209). Pifithrin-alpha-hydrobromide (Tocris biosciences, Cat # 1267).

### Histopathology

Lungs were perfused with 10 % formalin at 20 cm H_2_O and fixed overnight. Paraffin embedded tissues were sectioned at 4um thickness. For histopathology, tissues were subjected to haematoxylin & eosin staining and Masson’s trichrome staining.

### Immunofluorescence Staining

Tissues were deparaffinized in Xylene, rehydrated in ethanol gradient and permeabilized in 0.1 % triton X-100. Antigen retrieval was performed in citrate buffer at 95°C for 20 min. Sections were blocked (3 % BSA, 10 % donkey serum, PBS) and incubated with primary antibodies (SPC 1:1000 dilution, Scgb1a1 1:100 dilution, Scgb3a1 1:100 dilution, and Krt5 1:500 dilution) overnight at 4°C. Tissues were washed thrice and incubated with appropriate fluorescent secondary antibodies at 20°C, washed and mounted using prolong gold anti-fade mounting medium with DAPI (Life Technologies). Images were acquired on Zeiss microscope.

### Isolation of murine alveolar epithelial cells

Mice were sacrificed in the presence of isofluorane in a chamber. Lungs were exposed after bilateral thoracotomy and perfused with PBS containing 0.5 % EDTA. Lungs were digested in 20 mg/ml Dispase (Life Technologies, Cat # 17105-041) suspended in DMEM for 1 h at 37°C, then minced in presence of DNase I. The suspended cells were serial filtered through 100um, 70um and 40um. The cell suspension was stained with conjugated primary antibodies to PerCP-Cy5.5 anti-mouse CD45 antibody (Biolegend; Cat # 103132), APC-Epcam (eBioscience, Cat # 50-152-15) and dead cell stain sytox blue (Thermofisher, Cat # S34857). After final wash, cells were suspended in FACS buffer, analyzed by flow cytometry and sorted on BD Aria cell sorter for Epcam positive cells after CD45 exclusion.

### Isolation of human alveolar type 2 epithelial cells

Human lungs obtained from consented patients (IRB # 13-10738) at the time of transplant were perfused with heparin in PBS and digested with dispase (5U/mL) in DMEM for 30 min at 37^0^C in water bath. Lungs were minced and incubated in dispase solution for another 30 min at 37^0^C. DNase (0.25 mg/mL) was added to the lung digest and serial filtered through 70 𝜇m and 40 𝜇m filters. Lung homogenate was suspended slowly on 30 % percoll with 70 % percoll at the bottom of the tube. Centrifugation of lung homogenate and percoll gradient was carried out at 350 g for 20 min with no brake. The top layer of supernatant consisting of epithelial cells and immune cells was removed, to a new 50 mL tube and spun down at 350 g for 8 min with brakes. The supernatant was discarded and the pellet was washed 2X with PBS and resupended in DMEM supplemented with Penicillin, Streptomycin and Fungizone. IgG plates were prepared by coating petridishes with 0.5 mg/mL of human IgG in 50mM TRIS pH9.5 for 3 hours and washed 2X with PBS. Lung cell homogenate (5 mL) was added to each petridish and incubated for 60 min. The unattached cells were collected and spun down at 350 g for 5 min, washed in PBS twice followed by suspension in TEC+ media. Cells were counted again for post IgG panning numbers.

### Flow cytometry

Human lung cell suspension was stained with lysotrackerTM red DND-99 (Invitrogen, Cat # L7528) at 1:20,000 in a tissue culture incubator at 37^0^C for 45 min, washed in PBS, resuspended in FACS buffer and stained with conjugated primary antibodies to Alexa Fluor 647 anti-human CD326 EpCAM (Biolegend, Cat # 324212), pacific blue anti-human CD45 (Biolegend, Cat # 304022). Cells were washed 1X with PBS and resuspended in FACS buffer. Dead cell stain sytox blue was added prior to flow cytometry analysis. Cells were analyzed and sorted on BD Aria cell sorter. Data was analyzed on FlowJo V10.

### Three-dimensional organoid culture: Human lungs

Sorted CD45^-^Epcam^+^Lysotracker red^high^ cells (5000) were mixed with MRC5 fibroblasts (50,000) in 45 uL of TEC plus media. The cell suspension was mixed with 45 uL of Matrigel (Gibco) plated on the upper transwell chamber and allowed to solidify at 37^0^C for 10 min. To the bottom chamber of transwell, 500 uL of TEC plus media^44^ supplemented with 10uM of Y-27632 (ROCK inhibitor) was added. After 48 h in culture, the media in the bottom chamber was changed to TEC plus without Y-27632. Cultures were fed every 48 h and allowed to grow for 14 days at which time the images were captured.

### Mouse lungs

Sorted Epcam positive cells (5000) were mixed with NIH-3T3 fibroblasts (50,000) in 45 uL of TEC plus media. All the other steps for mouse lung organoid culture are similar to human lung organoid culture described above.

### TEC basic medium composition

1M Hepes 7.5ml, 200mM Glutamine 10ml, 7.5% NaHCO3 2 mL, Fungizone (1000X, 250 ug/ml) 500ul, Pencillin-streptomycin (100X, 10,000U or ug/ml) 5ml. Add DMEM/F-12 50:50 mix to a volume of 500 mL. The media prepared was sterile filtered, stored at 4^0^C and used within 4 weeks. TEC plus media composition: To 250 mL of TEC basic media, following reagents were added to make TEC plus. Insulin (10ug/ml) 250ul, Transferrin (5ug/ml) 250ul, Cholera Toxin (0.1ug/ml final) 25ul, Epidermal growth factor (25ng/ml) 25ul, Bovine Pituitary extract (15mg/500ml) 2.5ml, FBS (5% final) 12.5ml. The media prepared was sterile filtered, stored at 4^0^C and used within 2 weeks.

### Pre-processing scRNA-seq samples

Single-cell RNA-sequencing was performed on epithelial cell adhesion molecule (Epcam) positive cells from whole lung tissue of 5 mouse samples within 3 groups: 2 control TRF1^F/F^ mice, 1 early mouse and 2 late mice belonging to SPC-creTRF1^F/F^ genotype. The demultiplexed fastq files for these samples were aligned to the pre-built Cell Ranger mouse reference genome 2020-A using the 10x Genomics Cell Ranger v.6.1.1 count pipeline^45^ as described in the Cell Ranger documentation.

### Quality control of scRNA-seq data using Seurat

The filtered count matrices generated by the *Cell Ranger count* pipeline for all 5 samples were processed using the Seurat v3.1.4^46^ R package for single-cell RNA-seq analysis. Each sample was pre-processed as a Seurat object. For each Seurat object, the *Xist* gene was removed to avoid clustering by gender as Xist is present only in females. The top 1% of cells per sample with a high number of unique genes, cells with ≤200 unique genes and cells ≥10% mitochondrial genes were filtered out for each sample.

### Normalization and batch effects removal

The 5 samples were merged into a single Seurat object. Normalization and variance stabilization were performed using *SCTransform*^47^. Of the 5 samples, one control, one early timepoint and one late timepoint sample were sequenced using the single-cell 3’ v2 chemistry, while the remaining 2 were sequenced using the 3’ v3 chemistry. The sequencing chemistry was included as a variable to be regressed out for each feature during the SCTransform normalization. Based on the post normalization UMAP showing batch effects after regressing out the chemistry variable, we performed a second step using SCTransform (without adjusting for ‘sequencing chemistry’ variable), followed by *RunHarmony()* function that performs Harmony^48^ a tool for data integration and batch effects. Finally, we used RunUMAP with ‘harmony’ reduction and 1:20 dimensions to visualize the data using *DimPlot()* and *FeaturePlot()* Seurat functions. We grouped and split the UMAP based on the metadata to check for batch effects. No other batch effects were detected.

### Clustering

Clustering on the normalized Seurat object was performed using seven resolutions (0.2, 0.3, 0.4, 0.5, 0.6, 0.7, and 0.8) and 15, 20, 25, and 30 principal components (PCs). The Seurat function *FindNeighbors()* and *FindClusters()* was used to evaluate the UMAPs, and a heatmap reporting the number of clusters for each of the 28 settings (defined PC and resolution) was generated. PC=10 and resolution= 0.7 was selected corresponding to 28 clusters because at this resolution meaningful clusters were formed and no further granularity was observed beyond this resolution.

### FindMarkers for cell type annotation

To annotate the cell clusters, *FindMarkers()* function was used for each of the 28 clusters at PC=10 and resolution = 0.7 to obtain a list of cluster-specific gene markers expressed at logFC threshold = 0.1 and FDR = 0.01 using Wilcoxon Rank Sum test (as default).

### Subsetting the epithelial cell types

Based on the UMAPs and cell type annotation, the Seurat object was subsetted after normalization and harmony processing removing the non-epithelial cell types (clusters 7, 12, 19, 22, 23, 26). *SCTransform* and *Harmony* was applied on the subset of cells adjusting for the ‘*sequencing chemistry*’ variable. UMAPs for the three groups (control, early and late) were visualized. Seurat clustering and visualization was performed using DimPlot() as above and one setting (PC=10 and resolution = 0.6) was selected. Meaningful epithelial clusters were achieved at this resolution of 0.6 and no further granularity was observed beyond this resolution obtaining 27 clusters (Figure S1B). *FeaturePlot()* Seurat function was performed to visualize cells expressing markers of interest (Figures 1E, 2A, S2 and S3). Clusters with similar expression of specific marker genes were merged as evaluated using *FindMarkers()function*. Clusters 0, 1, 5, 18 were merged and annotated as AT2; clusters 6 and 10 as profibrotic AT2; clusters 4, 23 as Lyz1 High AT2; clusters 2, 7, 13, 22, 26 as Ciliated; clusters 3, 8, 11, 17 as Club; clusters 9, 12, 20, 25 as AT1 and clusters 14, 16 as Basal cells. Nine annotated epithelial cell types were obtained and 3 unknown cell types: clusters 15, 19, and 23 as reported in Figure 1C. The cell proportions were visualized across three conditions (control, early and late time-points) using *ggplot2* R package (Figure 1D). The function *VlnPlot()* was used to visualize the gene expression distribution of markers of interest across cell types (Figure 3A) and *DotPlot()* to visualize the average gene expression and the proportion of expression in specific cell types for markers of interest (Figures 3B, 3C and S5).

### Pseudobulking the counts and differential gene expression analysis

Differential gene expression analysis was performed by pseudobulking the counts of all the cells within each cell type. This analysis was performed on all the epithelial cell types. Clusters with low number of cells were removed, keeping clusters with a median of at least 5 cells across all individual biological replicates. To create the pseudocounts by cluster, *SingleCellExperiment* Bioconductor package^49^ was used to convert the Seurat object to single cell experiment object and *aggregateData()* function was applied and implemented in *muscat* R package (Multi-sample multi-group scRNA-seq data analysis tools). Then, the low count genes were filtered, keeping the gene with at least 5 counts in 2 or more samples. *DGEList()* was used and implemented in *edgeR* package to create the object including the filtered counts and the sample groups. A design was specified including the groups and the sequencing chemistry variable and the design was used when fitting the model using *glmQLFit()*. The *makeContrasts()* was used to create the 3 comparisons of interest: controls vs early timepoint; controls vs late timepoint and controls vs any timepoint. Each contrast was included in glmQLFTest() and up/downregulated genes were summarized using the *decideTestsDGE() and topTags()* to extract the results.

### Pseudotime Trajectory analysis

To identify lineage of cell differentiation, pseudotime or trajectory analysis was performed on the epithelial cells. First, the embedding of the cells from each cluster on the UMAP was evaluated. A subset of cells from some clusters were embedded on different area of the UMAP overlapping other clusters. For each cluster, the coordinates of the cells embedding were extracted and, after evaluating visually the outlier cells, the cells that were 1SD or 2SD or 3SD from the median of the UMAP coordinates for the given cluster were excluded. Then, the genes with low counts were filtered across cells. After cleaning the data, subsets of the Seurat object were created including combination of epithelial clusters of interest and trajectory analysis was performed using *slingshot* R package^23^ on 3 combination of subsets and starting cluster(Subset 1-starting cluster 0; subset 2 – starting cluster 0, and subset 1 starting cluster 3) to identify lineage of differentiation between cell states that was visualized on the UMAP (Figure 4A, 4C, S4A and S4C). Lineage 1 and 2 were selected for subset 1 – starting cluster 0; the unique lineage for subset 2 – starting cluster 0 and Lineage 2 and 3 for subset 1 starting cluster 3. The *tradeSeq (*version 1.10.0) R package^50^ was used to identify differentially expressed genes along the trajectory of interest. For each gene, tradeSeq fits a negative binomial generalized additive model (GAM) to smooth each gene’s expression in each lineage. Smoothers can be decomposed into a set of basis functions, which are joined together at knot points. The *evaluateK()* function was used on 100 genes to evaluate a range of knots (3-10) at 2 different seeds to be used when fitting the model. For the 3 subsets, 8-9, and 10 knots were selected. The fitGAM adjusting for ‘sequencing chemistry’ was performed with the selected number of knots and *associationTest()* and *startVsEndTest()* of the fitGAM object to extract potential changes in average gene expression along pseudotime. The function startVsEndTest uses a Wald test to assess the null hypothesis that the average expression at the starting point of the smoother (for eg. progenitor population) is equal to the average expression at the end point of the smoother (for eg. differentiated population). The patterns of the gene expression were visualized along the trajectories of interest with heatmaps showing the log2 transformed counts of a set of top 50-62 genes after p value adjustment with BH (Figure 4B, 4D, S4B and S4D).

### Pre-processing and library preparation for scRNA sequencing of human organoids

Organoids were digested with 0.25 % trypsin, centrifuged at 300g for 5 min. Cells were fixed according to the Parse Biosciences cell fixation protocol. Briefly, cells were resuspended in cell prefixation buffer followed by cell fixation. Fixed samples were resuspended in cell storage buffer and filtered through a 40 uM strainer and counted. Barcoding and library preparation was performed according to manufacturer guidelines in Parse Evercode WT Mini v3 single-cell Library preparation user manual version 1.3 – UM 0033 (Parse Biosciences). Each sample was loaded evenly, for a targeted amount of 5,000 cells per sample, and 2 identical sub-libraries of 5,000 cells each were prepared. Whole transcriptome sub-libraries were sequenced at the UCSF Cat Core on Illumina NovaSeqX sequencer on a 10B flow cell 300 cycle kit. For each sub-library 400 million reads were targeted producing about 100,000 mean reads per cell.

### Data analysis of Parse scRNA-seq data

The scRNA-seq dataset was processed, and analyzed using Trailmaker^TM^ (Parse Biosciences) pipeline module. The FASTQ files from each sub-library were demultiplexed. All sequence alignment and sample annotation steps were performed with Parse Biosciences split-pipe pipeline (v 1.1.0) in Parse’s Trailmaker Pipeline Module. Each Parse whole transcriptome sublibrary was aligned to the most recent human genome build (GRCh38), then aligned sublibraries were combined for analysis. Cells with <200 transcripts and the dead or dying cells with mitochondrial content >10 % were excluded. Doublets were filtered out using *scDblFinder* method. Data normalization, principal component analysis and data integration was performed using *Seurat* and *Harmony*. Clusters were identified using the *Leiden* method (resolution = 0.8) and a UMAP embedding was calculated to visualize the results. Cluster-specific markers were identified by comparing cells from one cluster to all other clusters. Cell type annotations were based on cluster specific markers. Feature plots and dot plots were generated to visualize cell-specific and differentially expressed genes.

### Statistics

Statistical analysis were performed with GraphPad Prism software (Version 10.0). P values were calculated using two-tailed Student’s t test. Welch’s correction was applied for analysis of unpaired groups. Differences in survival were analyzed by log-rank (Mantel-Cox) test. Data are represented as the mean ± SEM. P values represent the significance using the following symbols. ns = not significant, p>0.05, *p<0.05, **p<0.01, ***p<0.001, ****p<0.0001.

## Acknowledgements

This work was funded by the National Institute of Health HL RO1 139897 grant to Paul J. Wolters and UCSF Bakar Aging Research Institute Investigator Award to MB.

## Author Contributions

P.W. conceived the experiments, supervised the study and interpreted the data; R.N. designed and performed experiments, analyzed, interpreted data and wrote the manuscript; A.B., M.W. performed experiments. K.B. assisted with pathway analysis under the guidance of M.B., M.T., A.A, R.T. performed computational analysis on single cell data; J.K. contributed to acquisition of human samples; M.B. helped with interpretation of results and provided feedback. All authors approved final version of this manuscript.

## Supplemental information and figure legends

### Supplemental information

Document S1: Figures S1 – S8

## Supplementary Figures

**Figure S1.**
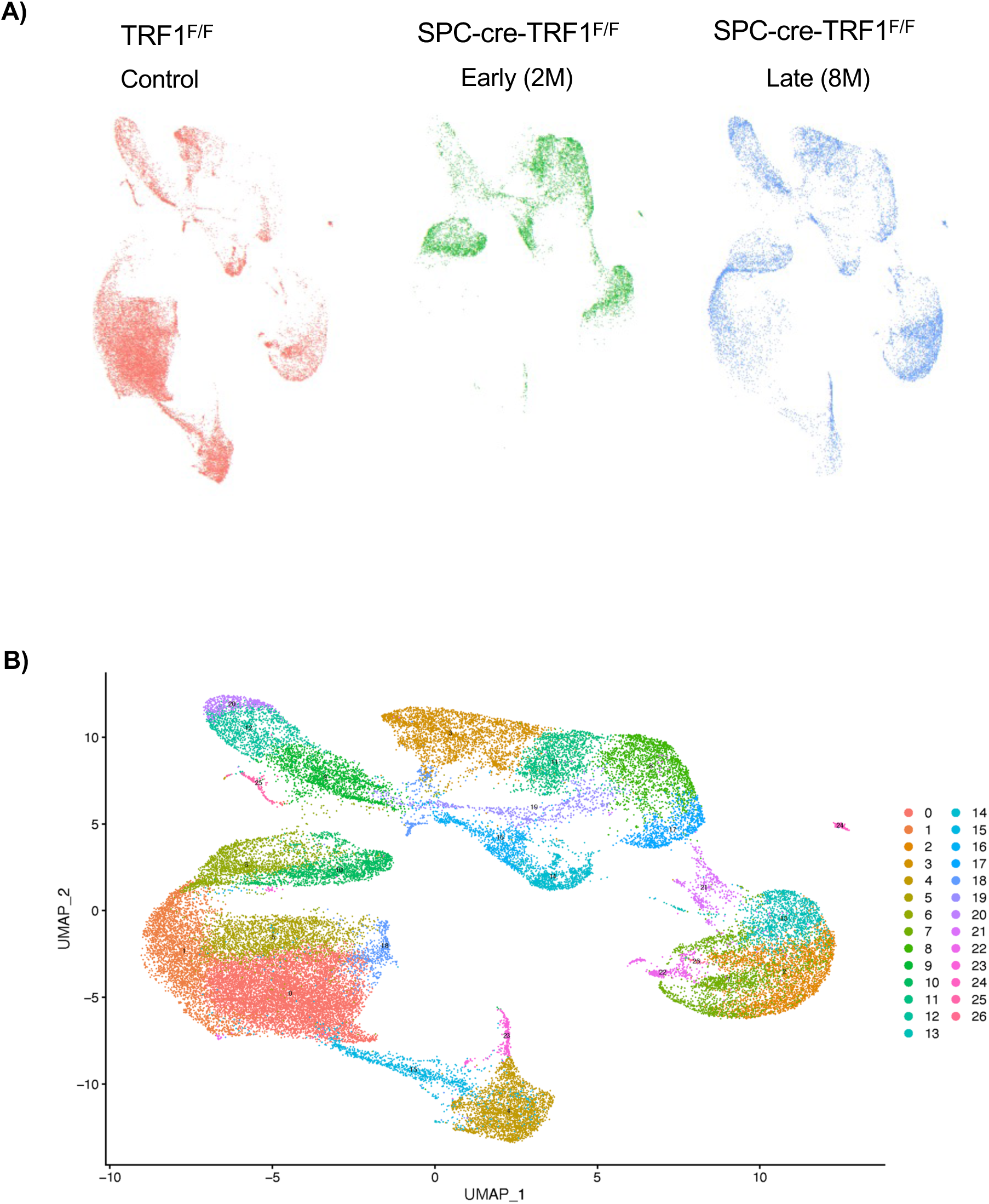
UMAP distribution of cells from single-cell RNA sequencing of mouse lung epithelium. **A)** UMAP plot displaying cell distribution in various clusters comparing control, early (2M) and late (8M) groups from TRF1^F/F^ controls and SPC-creTRF1^F/F^ mice with telomere dysfunction. **B)** UMAP plot indicating the total epithelial clusters from TRF1^F/F^ controls and SPC-creTRF1^F/F^ mice with telomere dysfunction.

**Figure S2.**
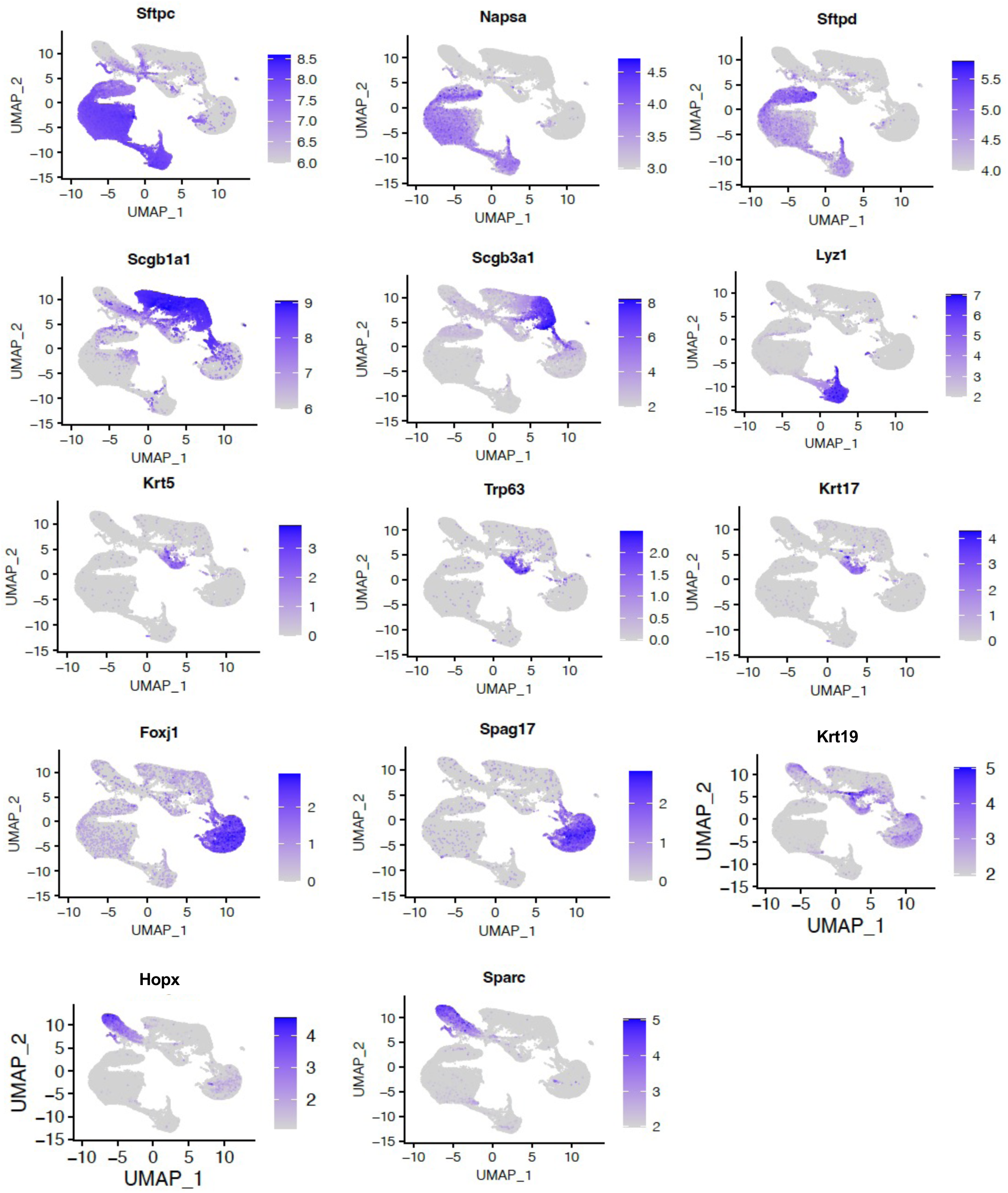
Markers indicating lung epithelial cell types. Feature plots showing expression levels of markers of lung epithelial types and sub-types. Color scale indicates fold expression. Blue=positive expression; Grey = No expression.

**Figure S3.**
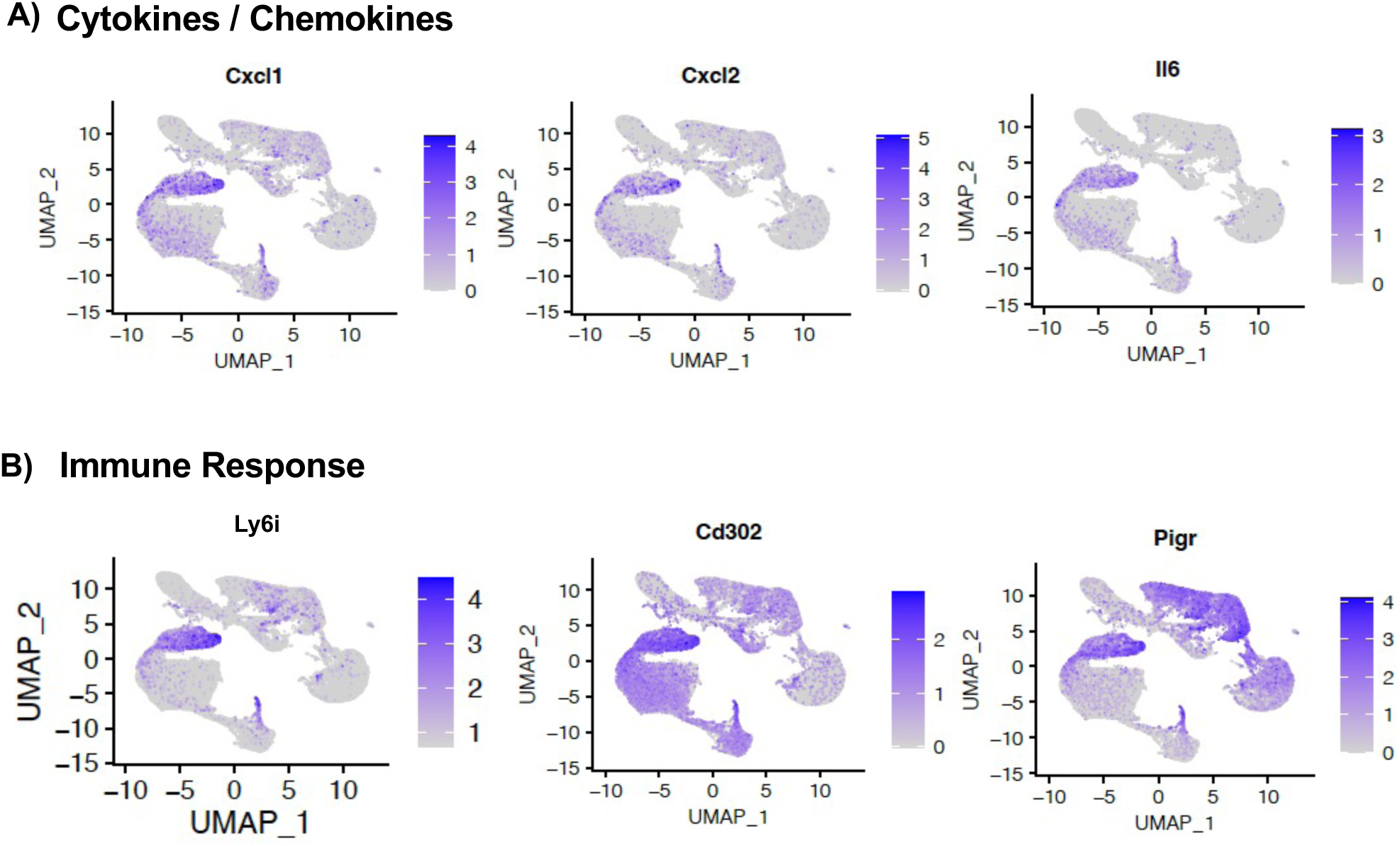
Markers indicating expression of cytokines, chemokines and markers of immune response in lung epithelial cell types. **A)** Feature plots showing expression levels of markers of cytokines and chemokines in lung epithelial clusters. Color scale indicates fold expression. Blue=positive expression; Grey = No expression. **B)** Feature plots showing expression levels of markers of immune response in lung epithelial clusters. Color scale indicates fold expression. Blue=positive expression; Grey = No expression.

**Figure S4.**
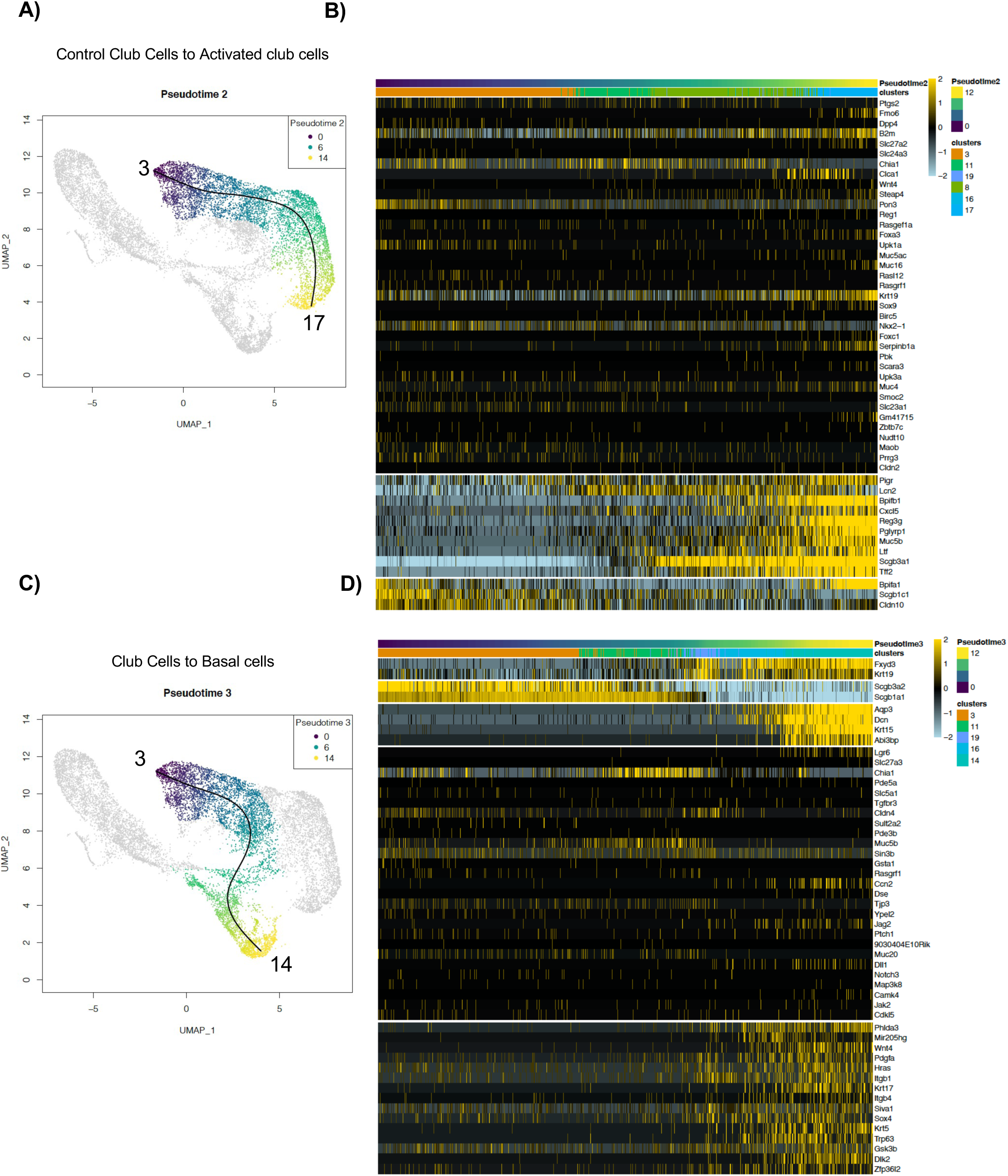
Trajectory of epithelial cells displaying the path from club cells to activated cell state. **A)** Slingshot trajectory showing the path of control club cells(purple) towards activated club cell clusters (yellow) (Pseudotime 2). **B)** Heatmap showing differentially expressed genes along the trajectory from cluster 3 to clusters 17. Color scale indicating relative expression from −2 (blue) to 2 (yellow). **C)** Slingshot trajectory showing the path of control club cells (purple) to basal cells (yellow) (Pseudotime 3). **D)** Heatmap showing differentially expressed genes along the trajectory from cluster 3 to 14. Color scale indicates relative expression from −2 (blue) to 2 (yellow).

**Figure S5.**
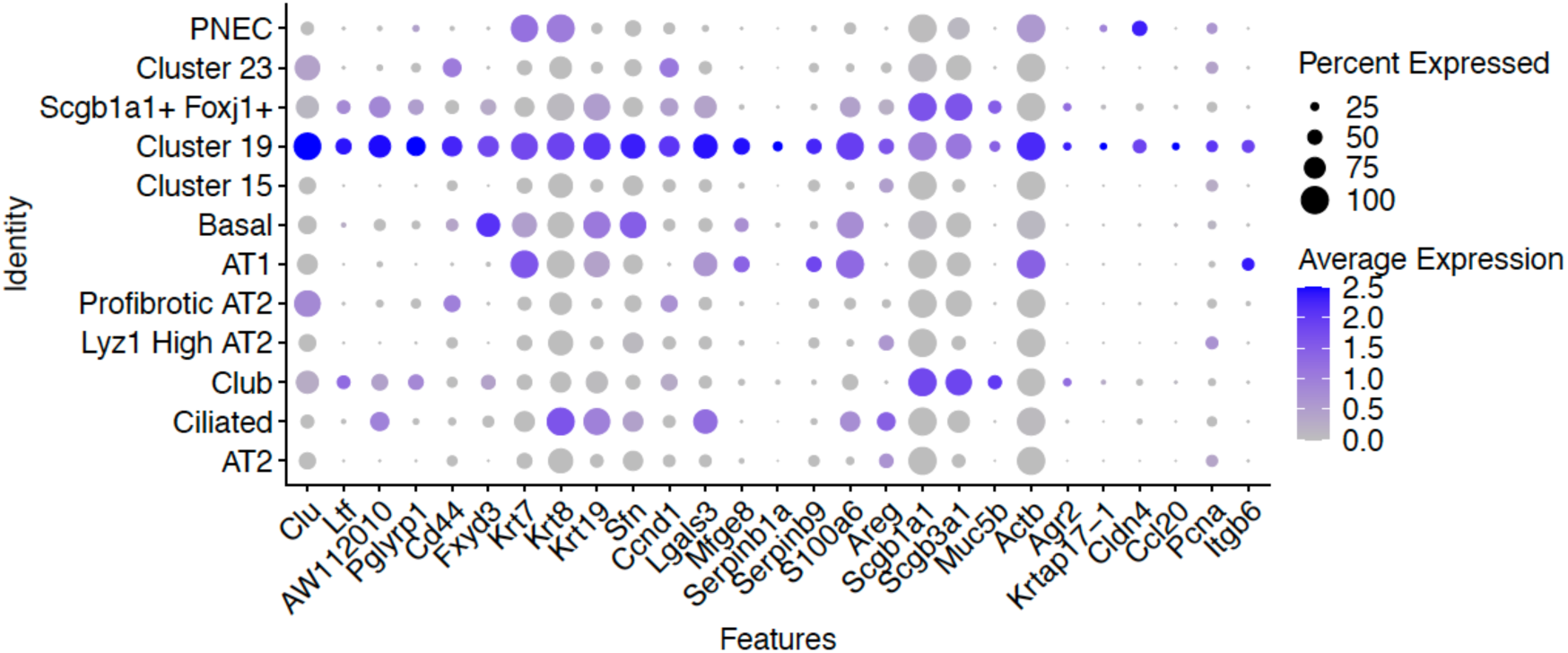
Dot plot showing cluster 19-specific gene expression. Dot plot showing average expression level of markers specific to cluster 19. Size of the dot indicates percent cells expressing the gene. Expression level is indicated by color. Blue=positive expression; Grey = No expression.

**Figure S6.**
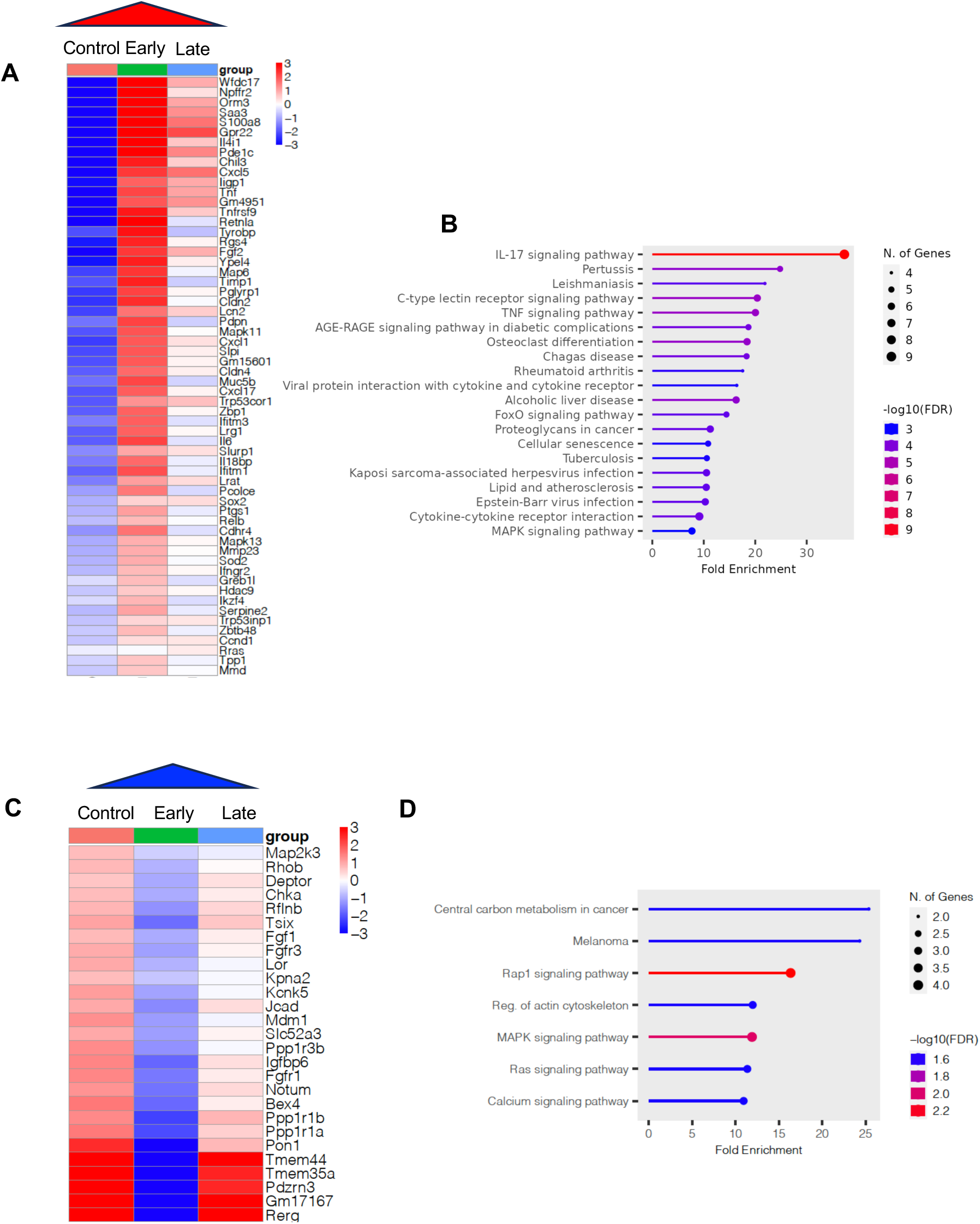
Differential gene expression and pathway analysis highlighting early timepoint changes. **A)** Heatmap of differentially expressed genes that are highest in early (red). **B)** KEGG pathway analysis showing pathway enrichment in response to upregulated genes in early. Size of the dot indicates number of genes in the dataset that overlap with the genes in the pathway. Color scale indicates enrichment FDR. **C)** Heatmap of differentially expressed genes that are lowest in early timepoint (blue). **D)** KEGG pathway analysis showing pathway enrichment due to down-regulated genes at early.

**Figure S7.**
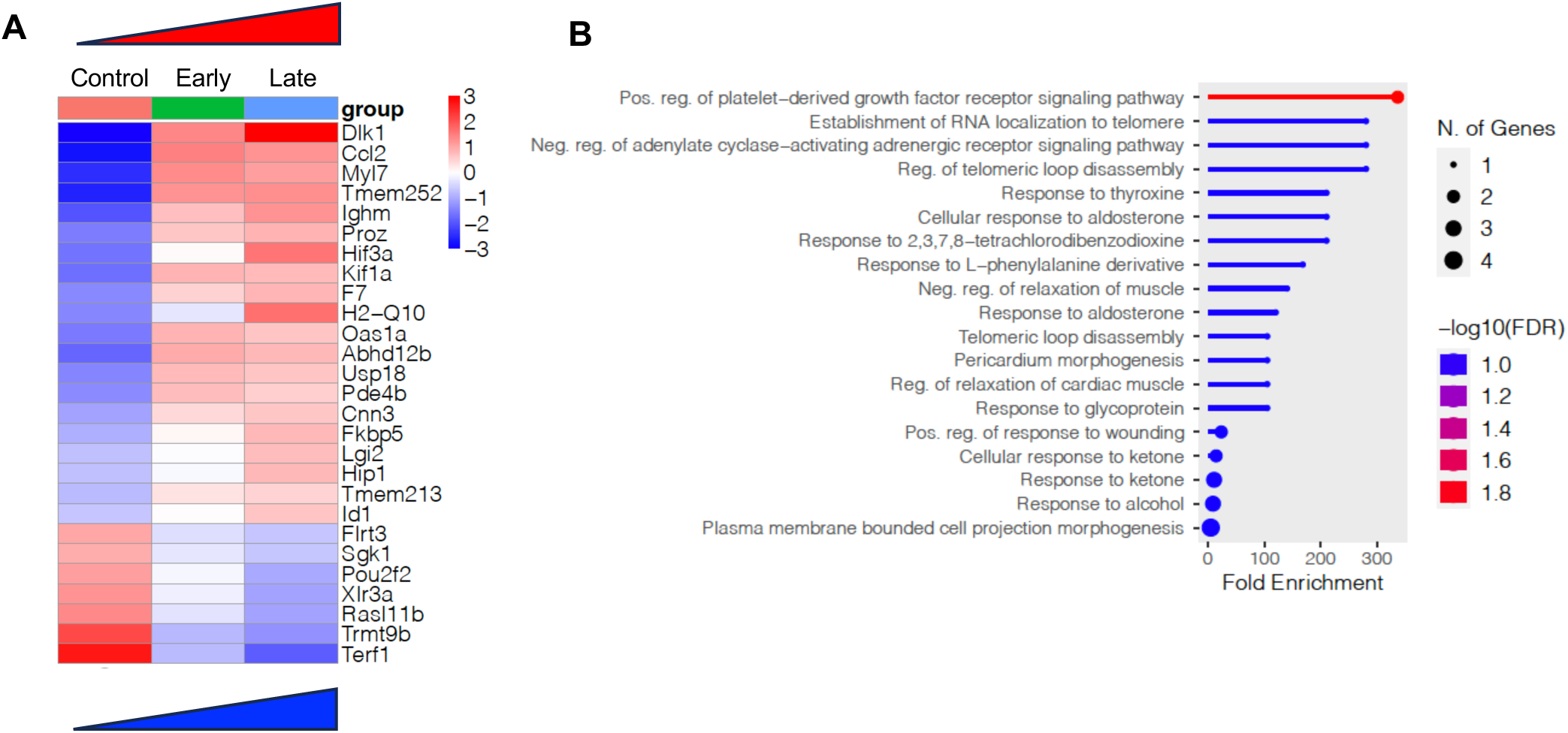
Differential gene expression and pathway analysis highlighting late timepoint changes. **A)** Heatmap comparing gene expression changes in control, early and late groups that are highest (red) and lowest in late group (blue). **B)** Gene Ontology (GO) biological process of differentially expressed genes that are highest (red) and lowest (blue) in late-timepoint.

